# Vitamin-B12 enrichment in tempeh by co-culture with *Propionibacterium freudenreichii* during fermentation

**DOI:** 10.1101/2022.11.06.515253

**Authors:** Meng Tian He, Kate S. Howell

## Abstract

Vitamin-B12, or cobalamin, is an essential human vitamin and most commonly acquired in the diet through consumption of animal products. Acquisition of the vitamin for people who follow strict plant-based diets is limited to supplementation via tablets or consumption of fortified foods. Tempeh — an Indonesian food of soybeans fermented with a filamentous fungi, *Rhizopus* sp. is a potential dietary source of cobalamin, but its presence is under question based on difficult quantification of this trace vitamin. This study tested the presence and concentration of vitamin-B12 in commercially available tempeh and tempeh produced with deliberate inoculation of various bacteria and moulds. Vitamin-B12 was not detected in any commercially available tempeh. Tempeh made with filamentous fungi and *Propionibacterium freudenreichii* (ATCC 9617) consistently produced detectable levels of cobalamin. *P. freudenreichii* co-fermented with *Rhizopus oligosporus* (ATCC 22959) produced the highest concentration of cobalamin at 8.26 ± 0.13 μg/100g of wet weight of tempeh. Co-fermentation with a different tempeh mould significantly reduced the amount of cobalamin (*P* = 0.008). Results from this study suggest that incorporating nutrient-enhancing microbes into an existing fermentation stage of a product is an effective method to increase the nutritional density of food. The results of this study suggest that cheap, safe and easily cultured microbes can contribute to the nutritional diversity of people following plant-only diets.

## Introduction

Cobalamins are a group of structurally similar, cobalt-ligating molecules which are commonly referred to as vitamin-B12 and are essential for numerous functions for human health (Kennedy & Taga, 2020). Although it is understood that all naturally sourced cobalamin is initially synthesised by bacteria, cobalamin accumulates in animals and thus makes animal products the most abundant dietary sources of the vitamin (Watanabe & Bito, 2017). Increasing prevalence of veganism and vegetarianism (Jones, 2020) has piqued scientific interest in plant-based sources of vitamin-B12 to cater for people following these dietary rules (Watanabe, Yabuta, Bito, & Teng, 2014). Initial research has found traces of cobalamin in seaweed, spirulina, and soy products (Herbert & Drivas, 1982; Truesdell, Green, & Acosta, 1987; van den Berg, Dagnelie, & van Staveren, 1988). However, studies critical of the quantification methods have called for further clarification of the issue (Dagnelie, van Staveren, & van den Berg, 1991).

Early investigation into tempeh — a traditional food from Indonesia where soybeans are fermented with *Rhizopus* mould (Hartanti, Rahayu, & Hidayat, 2015) — measured vitamin-B12 activity in samples containing gram-negative, rod-shaped bacteria (Liem, Steinkraus, & Cronk, 1977). These bacteria were identified as *Klebsiella pneumoniae* (Truesdell et al., 1987) and *Citrobacter freundii* and the responsible microbes for producing vitamin-B12 in tempeh (Keuth & Bisping, 1993). Additionally, production of cobalaminn by *C. freundii* was two times higher than when fermented with *Rhizopus oligosporus* than a control of the bacterium alone, implying that the main process of tempeh production itself helps to enhance vitamin-B12 production in these bacteria. Some researchers have focused on using generally-recognised-as-safe (GRAS) vitamin-B12 producing bacteria in lupin-based, tempeh, and have successfully augmented its vitamin-B12 content (Wolkers-Rooijackers, Endika, & Smid, 2018), but has not been attempted in soybean-based tempeh. Considering all these factors in conjunction with associated implications for sustainable food production, researchers have highlighted the potential for tempeh as a cheap, nutritious food, particularly as a source of vitamin-B12 (Ahnan-Winarno, Cordeiro, Winarno, Gibbons, & Xiao, 2021; Tamang, Anupma, & Nakibapher Jones Shangpliang, 2022).

Adequate detection and quantification of cobalamins in food has proven difficult; microbiological assays commonly employed have been deemed inadequate, as these assays also detect nutritionally inactive cobalamin analogues (Degnan, Taga, & Goodman, 2014). However, new methods of vitamin-B12 quantification in foods have been devised (Chamlagain, Edelmann, Kariluoto, Ollilainen, & Piironen, 2015; Okamoto, Hamaguchi, et al., 2020). Here, we use a verified assay to quantify the cobalamins present in commercially available tempeh, and then use this same assay to test the cobalamin in tempeh made with deliberate introduction of different bacteria. We show that introduction of *P. freudenreichii* in tempeh fermentation can produce appreciable and nutritionally relevant amounts of vitamin-B12, in a sensorially-attractive food product. Inclusion of GRAS-cobalamin producers into the tempeh fermentation process provides an harmonious way of producing a biofortified plant-based food. We suggest this method to complement the many plant-based products on the market that are commonly fortified with cobalamin, isolated from a bacterial source (Damayanti et al., 2018).

## Materials and Methods

### Sample preparation for measurement of vitamin-B12

Five different commercially available tempeh were purchased from supermarkets and markets in Melbourne Australia; Macro Tempeh, Nature’s Kitchen, Nutrisoy, Tally Ho Farms, and Primasoy. All tempeh samples were homogenised using a hand blender to create a paste, and were weighed. This paste was then spread approximately 1 cm thick onto trays, and frozen overnight. Samples were freeze dried for roughly 96 hours and weighed again to establish the moisture content of the tempeh samples. The samples were then stored until analysis in dark, covered containers at -20°C to minimise chemical activity and cobalamin degradation.

### Chemicals and standards

All chemicals were purchased from Sigma-Alrich unless otherwise specified. Sodium acetate buffer stock was made by dissolving 4.1 grams of anhydrous sodium acetate in water and adjusted to pH 4.5 using anhydrous acetic acid. Cyanide salt stocks of 1% weight to volume (w/v) were prepared using KCN and water.

### Sample Preparation

Freeze dried tempeh paste samples were ground into a fine powder using a mortar and pestle; 25 g of each sample was then dissolved and vigorously mixed with 100 mL of water to make homogenous tempeh solutions and analysed by the method outlined for Easi-Extract® Vitamin-B12 (R-Biopharm, 2020)..

Liquid samples were weighed into aliquots and 0.05g of Taka-Diastase from *Aspergillus oryzae* (Sigma-Aldrich) in 1 mL of 1% w/v KCN solution, and mixed. Samples were incubated in a 40°C water bath for 30 minutes with gentle shaking. Subsequently, 50 mL of sodium acetate buffer was added and mixed, and samples were then heated in a 100°C water bath for 30 minutes with shaking. After cooling, solutions were transferred to 100 mL volumetric flasks, and made up to 100 mL with MilliQ water. These solutions were filtered through Whatman 1 filter paper, and then through 0.22 µm syringe filters (Thomas Scientific).

The filtrates were added to pre-prepared Easi-Extract® Vitamin-B12 (LGE) immunoaffinity columns, and prepared in accordance with the manufacturer’s instructions (r-Biopharm, 2020). The immunoaffinity column facilitated isolation and concentration of cyanocobalamin in samples using monoclonal antibodies of high binding specificity. The solution from the column was dissolved in 0.1% formic acid, added to fused-insert macro vials (ThermoFisher; Sydney Australia) capped and tested using HPLC.

### High Performance Liquid Chromatography

HPLC quantification was conducted using a reverse-phase Ascentis® Express RP-Amide, 5µL column (Supelco) and UV detection with an Agilent 1260 Infinity II equipped with a flexible pump and DAD detection using a wavelength of 361 nm. OpenLab CDS was used to record and analyse the data. Injection volumes were 5µL, and the flow rate of the systems were set to 0.5 mL/min. Two mobile phases were required for HPLC analysis. Solution A was 0.1% formic acid (Thermo Fisher) by volume in water, and Solution B was 0.1% formic acid in methanol (Thermo Fisher). Each buffer was degassed using ultrasonication. Analytical-grade _≥_98% cyanocobalamin (Sigma Aldrich; V2876) was dissolved in MilliQ water standard (Sigma Aldrich; V2876) and used to create a standard curve of peak area and concentration. The gradient conditions used for HPLC were 70% of Solution A at 0 minutes, to 65% at 10 minutes, to 70% at 14 minutes, and 70% at 17 minutes.

### Microbial growth conditions

Bacteria and mould isolates were sourced from ATCC (Table 1). Different broth, agar and incubation conditions were used to support the growth of each isolate (Table 1). Mould and bacteria were isolated from an unpasteurised commercial tempeh (Primasoy; Victoria Australia) using spread plate and dilution series. Here, one gram of the tempeh was homogenised in sterile water to create a solution for serial dilution. Duplicate spread plates were made with dilutions on plate count agar (PCA; Oxoid, Sydney, Australia) and incubated at 28°C for 48 hours. Colonies of different morphology were selected and subcultured until purified on PCA. Single colonies of bacteria were identified using 16S rRNA primers using a commercial service provided by the Australian Genome Research Facility (St. Lucia, Queensland, Australia) and the results deposited in an online sequence database (NCBI-here).

**Table 1:** Organisms used in this study, with the media and incubation conditions used to create their concentrated pure cultures. Unknown moulds and bacteria were isolated from a sample of the same unpasteurised, commercial tempeh.

| Organism | Abbreviation | Media | Source | Conditions |
| --- | --- | --- | --- | --- |
| Moulds |  |  |  |  |
| *Rhizopus oligosporus* | RO | Potato Dextrose Agar (PDA; Oxoid) | Commercial starter | 28°C, Aerobic |
| Unknown mould | PS |  | This study |  |
| Bacteria |  |  |  |  |
| *Propionibacterium freudenreichii* subsp. *freudenreichii* | PF | Yeast extract lactate agar | (Malik, Reinbold, & Vedamuthu, 1968) | 28°C, Anaerobic |
| *Citrobacter freundii* | CF | Nutrient Agar (Oxoid) | ATCC | 37°C, Aerobic |
| *Lactobacillus plantarum* | LP | De Mann, Rogosa, and Sharpe (MRS) agar (Oxoid) | (Winters et al 2019) | 28°C, Anaerobic |
| *Fructilactobacillus sanfranciscensis* | LS |  |  |  |
| *Lactobacillus rhamnosus* (LGG) | LGG |  |  |  |
| *Lactobacillus casei* | LC |  |  |  |
| *Bifidobacterium animalis* (BB12) | BB12 | 0.05% L-cysteine MRS agar | (Laxmi, Mutamed, & Nagendra, 2011) | 28°C, Anaerobic |
|  |  | (Sigma Aldrich, C7352) |  |  |
| *Bacillus* sp. | B1 | Tryptic soy agar (Oxoid) | This study | 28°C, Aerobic |
| *Leuconostoc citreum* | B2 | MRS agar | This study | 28°C, Anaerobic |

### Experimental tempeh production

Dried food-grade soybeans (Primasoy; Victoria, Australia) were soaked for 24 hours, drained and steamed for 30 minutes. The steamed soybeans were soaked for a further 20 hours, dehulled, drained, and autoclaved at 121°C for 30 minutes, then drained and cooled before microbial inoculation and incubation. Tempeh was made using sterilised soybeans, enclosed in plastic bags with 1 mm perforations spaced 1.5 cm apart from each other. Beans were inoculated using 10^−4^ CFU/g of isolated spores suspended in sterile water and glycerol, and 10^−7^ CFU/g of concentrated bacterial broth cultures.

Treatment samples were made using one of the two moulds and one of nine bacteria to make 18 treatment samples in total. Two batches were made with one of each of the moulds only, as negative controls for bacterial action. Uninoculated soybeans were also incubated as a non-fermented control to produce 21 different samples in total. Each fermentation and control treatment was done in duplicate. Samples were taken for analysis of vitamin-B12 production as detailed above.

### Statistical Analysis

Two tailed t-tests were conducted to test significant differences between different treatments (_D_ = 0.05) using Microsoft Excel and the Analytic Solver® Data Mining add-on (Microsoft Corporation, Redmond, Washington, USA).

## Results and discussion

Soybean-based tempeh is a nutritious plant-based food product used as a traditional food in Indonesia and widely available commercially as a protein-rich dietary addition, especially to those people following a plant-based diet. To investigate the potential of tempeh to be a dietary source of vitamin-B12, commercially available tempeh were analysed using verified analytical methods by HPLC. Tempeh was also made using various combinations of bacteria and moulds to develop robust fermentation protocols with alternative starter cultures to fortify vitamin-B12 in tempeh.

### Vitamin-B12 was not detected in commercially available tempeh

HPLC analysis of commercial tempeh samples showed that none contained detectable traces of cobalamin (Supplementary Data). Vitamin-B12 was long found to be present in tempeh (Liem, Steinkraus, & Cronk, 1977; Keuth & Bisping, 1993), yet these studies used microbial methods of quantification that have been shown to have cross-reactivity with molecules that are not biologically active in humans as vitamin-B12, including pseudovitamin-B12, DNA, deoxyribonucleotides, and deoxyribonucleosides (Sauberlich, 1999). Of the samples tested here, commercially available tempeh may have been pasteurised and it is possible that vitamin-B12 is not heat labile under these conditions. Yet, the Primasoy tempeh tested was known to be unpasteurised, did not show detected vitamin-B12 and was selected for further microbial analysis.

### Unpasteurised tempeh contains two bacterial types

We hoped to find microbes in the commercial preparation of tempeh that had cobalamin-synthesising activities. We isolated two distinct bacterial colonies from the Primasoy tempeh, and found a fragment of 700 base pairs of amplified regions from the 16S rRNA best matched an unidentified *Bacillus* sp. and *Leuconostoc citreum*.

The *Bacillus* isolate was present in concentrations of roughly 10 CFU/g of tempeh. Low levels of the *Bacillus* spp. could be attributed to acidification of soybeans during the soaking step of tempeh, as *Bacillus* growth can be controlled for by this method. For example, acidification of soybeans beans to a pH below 5 was able to successfully prevent *Bacillus cereus* growth in tempeh in some studies (Nout, Beernink, & Bonants-van Laarhoven, 1987). *Bacillus* bacteria are commonly found in soil, and some studies have shown that some particular species are endemic to the soybean rhizosphere (Wahyudi, Prasojo, & Mubarik, 2010). As such, their presence in tempeh may be a result of risk factors associated with soybean production and harvest. *B. cereus* has also been specifically identified as a relatively common contaminant in tempeh (Keuth & Bisping, 1994).

The second bacterium identified, *Leuconostoc citreum*. is gram positive, rod-shaped, GRAS, lactic-acid bacterium, often associated with fermented foods that, notably, has the capacity to reduce the pH (Hemme & Foucaud-Scheunemann, 2004). *Leuconostoc* sp. are often considered health promoting probiotics, and desirable in most foods (Soomro, Masud, & Anwaar, 2002). These bacteria were found in low concentrations, at roughly 95 CFU/g of tempeh. This may be a result of the production of antimicrobial compounds of the tempeh mould (Ito et al., 2020; Kobayasi, Okazaki, & Koseki, 1992; Wang, Ruttle, & Hesseltine, 1969).

Experimental tempeh made using *Rhizopus oligosporus* and either *Bacillus* spp. B1 and *Leuconostoc citreum*. B2 did not have detectable levels of vitamin-B12 (Supplementary figure 1). Thus, microbial members of the fermentation ecosystem do not produce detectable levels of vitamin-B12 in the context of commercial tempeh production, which was reflected in the results of the analysis of commercial tempeh samples. This may be because these species do not produce vitamin-B12 in general, or their concentrations, the microbial community, or tempeh production conditions are not conducive to cobalamin production.

### Tempeh fermented with *P. freudenreichii* and *F. sanfransiscensis* contains vitamin-B12

Of the tempeh made with deliberate inoculation of Rhizopus mould with bacteria (Table 1), the formulations with *P. freudenreichii* were the only ones that consistently produced detectable levels of cobalamin; one replicate of *F. sanfransiscensis* fermented with *R. oligosporus* also managed to yield a detectable concentration of cobalamin (Table 2).

**Table 2:** Bacteria and the amount of detectable cobalamin produced at 28°C in a soybean substrate for 48 hours, as tempeh. Expressed in their relation to CFU and mass, as their mean values ± standard deviation. * = single replicates. ND = Not detectable.

| Bacteria | Mould | Average amount of B12 Produced ($\mu\text{g}/10^7$ CFU) | Average B12 in Tempeh ($\mu\text{g}/100\text{g}$) |
| --- | --- | --- | --- |
| *P. freudenreichii* | *R. oligosporus* | $0.083 \pm 0.0013$ | $8.264 \pm 0.13$ |
| | PS Mould | $0.012 \pm 0.0089$ | $1.188 \pm 0.89$ |
| *F. sanfransiscensis* * | *R. oligosporus* * | $0.0068$ * | $0.681$ * |
|  | PS Mould | ND | ND |

*P. freudenreichii* is used in industrial production of vitamin-B12, presumably for its cobalamin synthesising efficiency (Martens, Barg, Warren, & Jahn, 2002). It has often been used in many cobalamin bio-enrichment studies to successfully augment foods with vitamin-B12 (Xie et al., 2018; Xie et al., 2019). Furthermore, it is GRAS, and has been demonstrated to have the capacity to produce particularly high quantities of active forms of cobalamin rather than pseudocobalamin (Deptula et al., 2015).

The formulation producing the most vitamin-B12 was PF+RO, which, on average, contained 8.26 ± 0.13 μg of cobalamin per 100 g of tempeh. When paired with the commercial mould, *P. freudenreichii* seemed to synthesise significantly less (*P* = .008) cobalamin than the PF+RO sample (Table 2). This suggests that there is an interaction between the bacteria and mould affecting the final quantity of detectable vitamin-B12 in tempeh made in this manner. Unfortunately, based on the experimental design of this study, it is not possible to confidently assert why or how this interaction occurs. This phenomenon could be due to the two species of mould having different antimicrobial effects (Ito et al., 2020; Kobayasi, Okazaki, & Koseki, 1992; Wang, Ruttle, & Hesseltine, 1969), or perhaps they may produce different metabolites and by-products during fermentation that may affect cobalamin concentration directly. Conversely, some studies contend that *R. oligosporus* can have a cobalamin enhancing effect when fermented with *P. freudenreichii* (Signorini et al., 2018). Therefore, the difference between the two moulds could be attributed to different magnitudes of this same effect. Alternatively, a combination of an inhibitory effect from PS and an augmenting effect from the RO may also explain this difference.

Having already been successfully used to biofortify various cereal products with cobalamin (Xie et al., 2018; Xie et al., 2019), *P. freudenreichii* is an exceptional candidate for vitamin-B12 biofortification in general. Due to the ease of which novel microorganisms can be incorporated into their production systems, fermented foods seem to be particularly suited for these biofortifications. In fact, the unique effects of microbial growth is one of the main reasons behind why fermented foods are considered to be healthy and beneficial. For example, researchers have highlighted how various fermentation processes can improve the bioavailability of minerals in plant-based foods (Samtiya, Aluko, Puniya, & Dhewa, 2021). It only makes sense to further study the potential for fermentation to enhance nutritional density of foods — whether that be from altering substrates, temperature, atmospheric conditions, or microbial cultures.

One replicate of tempeh fermented with *F. sanfranciscensis (*LS+RO) contained 0.425 μg/100g of vitamin-B12. The cobalamin-producing capacity of *F. sanfranciscensis* has not been extensively reported. Vogel et al. (2011) found that eight out of eleven strains of *F. sanfranciscensis* were able to grow on vitamin-B12 free media and use this to claim that they can produce cobalamin *de novo*, despite only having one gene involved in vitamin-B12 synthesis. However, this thinking is flawed, as several bacterial species have been shown to be able to grow without supplementary cobalamin. For example, *Lactobacillus delbrueckii* subsp. *lactis* (ATCC ® 7830™) — the bacteria whose growth is used to quantify cobalamin in microbial assays — can substitute cobalamin with deoxyribosides, deoxynucleotides, or pseudocobalamin, and can therefore grow in the absence of vitamin-B12 in some situations (Watanabe & Bito, 2018). Having only one gene in cobalamin synthesis suggests that *F. sanfranciscensis* is able to partially synthesise the molecule, a relatively common phenomenon in various bacteria. Sokolovskaya, Shelton, and Taga (2020) highlighted the interactions of non-cobalamin cobamide-producing microorganisms, and mentions several bacteria which are able to produce cobalamin-like molecules. The data presented here would suggest that production of cobalamin in tempeh is not possible by this particular species of *F. sanfranciscensis* by itself; however, incorporating a more complex community of microorganisms may facilitate cobalamin biosynthesis.

### Biofortified tempeh as a rich source of vitamin-B12

Tempeh is often sought as a nutritionally dense food for those people following a strict plant-based diet due to it containing vitamin-B12. Our study shows that commercial tempeh in Australia does not contain vitamin-B12, and note that none of the commercial preparations here claim to contain vitamin-B12. It is possible that increasing sterility in tempeh production, due to food safety concerns and regulations, preclude the unintentional activity of bacteria that were thought to be sources of cobalamin in the fermentation of tempeh in the past. Here, we suggest the potential for a cobalamin-producing bacteria to be incorporated into commercial tempeh production, to produce both appealing, nutritionally rich, and achievable within existing tempeh production lines.

Results from this study indicate that *P. freudenreichii* would be a good inclusion in a tempeh mixed culture for the purposes of vitamin-B12 enhancement, resulting in a visually-attractive tempeh (Supplementary figure 2). The results presented here show that tempeh fermented with *R. oligosporus* and *P. freudenreichii* biofortified cobalamin content to 8.264 ± 0.13 µg/100g of tempeh. Consumption of just 10g of tempeh would result in consumers exceeding the recommended daily intake (RDI) of 2.4 μg of vitamin-B12 for adults suggested by the Food and Nutrition Board of the Institute of Medicine (1998), and the Australian Government National Health and Medical Research Council (2018). While studies have shown that the average Australian consumer’s daily vitamin-B12 intake exceeds this amount at 5.5 and 4.7μg for men and women respectively, others have noted that vegan populations have particularly low blood serum levels of vitamin-B12 when compared to the average omnivore (Truswell, 2007).

The method of analysis using a commercial immunoaffinity column with HPLC is specifically targeted to the bioavailable structural variants of cobalamin. However, the specific bioavailability of cobalamin from tempeh produced in this manner is unknown. As such, analogous compounds may still be present in these samples. This is significant because analogues may have the potential to inhibit vitamin-B12 absorption, and could therefore have anti-nutritive effects (Zeuschner et al., 2013). This was the case for spirulina, once touted as a high source of plant based vitamin-B12 — it is now known that it is actually high in pseudocobalamin which can inhibit vitamin-B12 absorption (Dagnelie et al., 1991; Herbert & Drivas, 1982). Therefore, further research into other corrinoids produced by the method of fermentation explored in this study would be required to determine the nutritional relevance of these results. Additionally, to investigate the validity of vitamin-B12-biofortified tempeh in a consumer context, further studies should also be conducted on the effects of food processing after fermentation, such as pasteurisation, cooking, and storage to validate stability and availability of vitamin-B12 for consumers.

Finally, it should be noted that the sensory properties of tempeh can significantly change based on differences in its fermentation parameters. Beyond appearance and texture of the tempeh cake that is formed, the flavour of the product can be affected by variations in chemical concentrations due to different microbial metabolites (Wikandari et al., 2021). As such. further research is required to investigate the flavour changes in tempeh fortified with vitamin B12 producing bacteria like *P. freudenreichii*.

### Conclusion

Cobalamin is an essential vitamin for human health; it needs to be acquired externally as it can not be synthesised by humans, and without it serious disease and death may occur. People who consume little or no animal products, such as vegans, are particularly at risk of deficiency because it is typically acquired from ingesting meat or dairy products. Tempeh, a high protein, plant-based fermented food, has been reported to be a source of vitamin-B12 due to the presence of bacterial contaminants, and could be a practical means of providing dietary vitamin-B12 in these particular populations.

Commercially available tempeh in Australia were found to contain no detectable levels of cobalamin. This may be due to pasteurisation, starter culture usage, and processes that minimise bacterial contamination and thus, inactivation or depletion of this essential vitamin. We isolated two bacteria from an unpasteurised commercial tempeh which were identified as *Bacillus* spp. and *Leuconostoc* spp. Tempeh made with supplementation of these bacteria also did not show detectable vitamin-B12.

Tempeh biofortification with vitamin-B12 was achieved by incorporating a vitamin-B12-producing bacteria, *P. freudenreichii* subsp. *freudenreichii* (ATCC 9617) into an existing fermentation step. The amount of cobalamin produced in tempeh by this method was found to be significantly affected by the type of mould that it was co-fermented with (*P* = .008). *P. freudenreichii* fermented with *R. oligosporus* produced tempeh with the highest concentration of vitamin-B12, yielding a concentration of 8.264 ± 0.13 µg/100g wet weight. Consumption of 100g of tempeh would more than adequately meet the recommended dietary intake for vitamin-B12. We recommend deliberate inoculation of *P. freuendreichii* into tempeh fermentation to make nutritionally dense, sensorially attractive tempeh.

## Credit Statement

Meng Tian He: Conceptualization (lead); writing – original draft (lead); formal analysis (lead); writing – review and editing (equal); Kate Howell: Conceptualization (supporting); Writing – original draft (supporting); Writing – review and editing (equal)

## Acknowledgements

Dr Hafiz Suleria is thanked for useful discussions for analytical method construction. Xinwei Ruan is thanked for her contributions to uploading sequences to Genbank.

## Data Availability statement

The datasets generated and/or analysed during the current study are available in the NCBI repository at *[persistent link to datasets]* or are included in this published article.

## Supplementary figures

**Supplementary figure 1:**
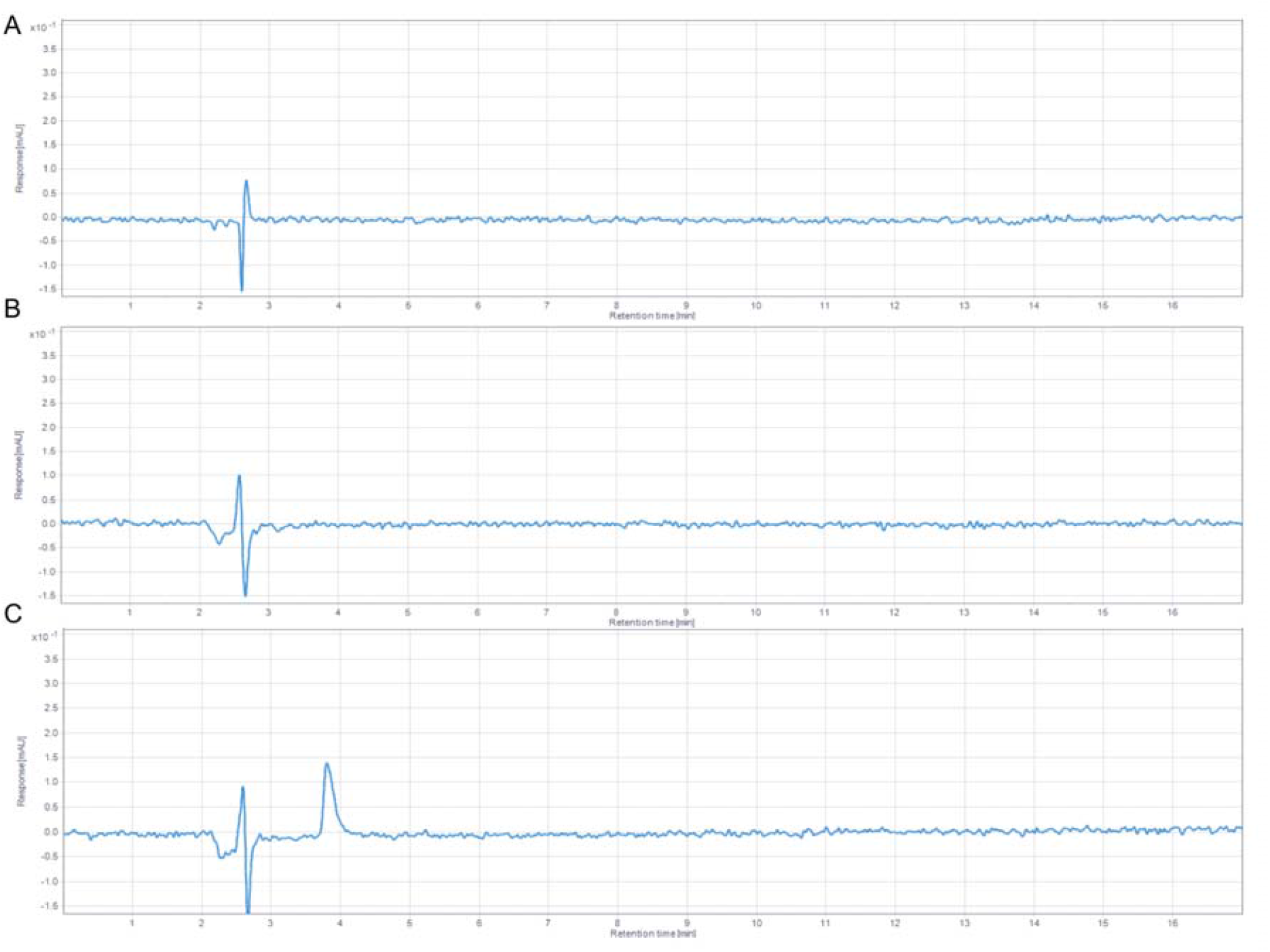
Example chromatograms of: (A) a tempeh sample inoculated with *Bacillus* sp. B1, (B) a tempeh sample inoculated with *L. citreum* B2, and (C) a spiked vitamin-B12 containing sample.

**Supplementary figure 2.**
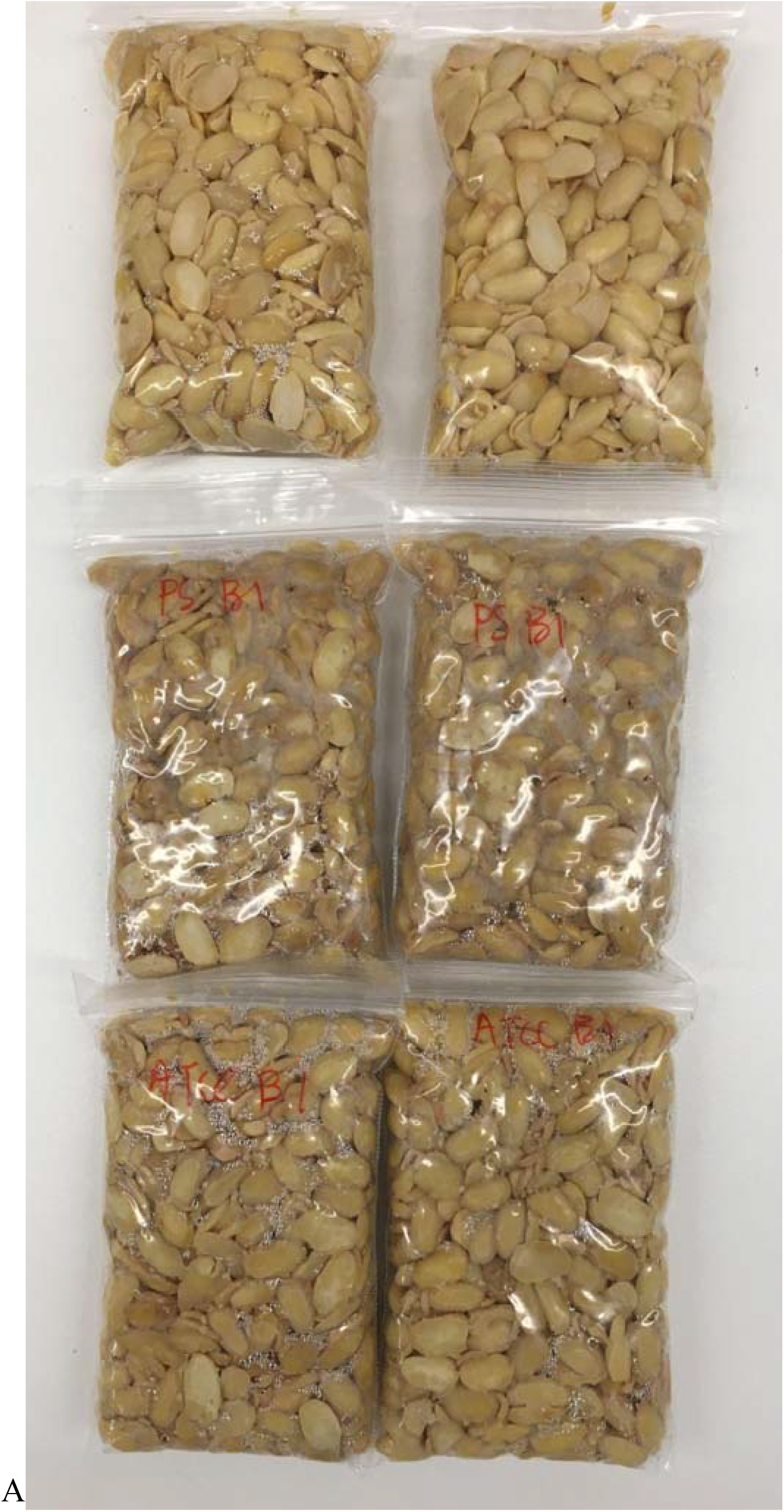
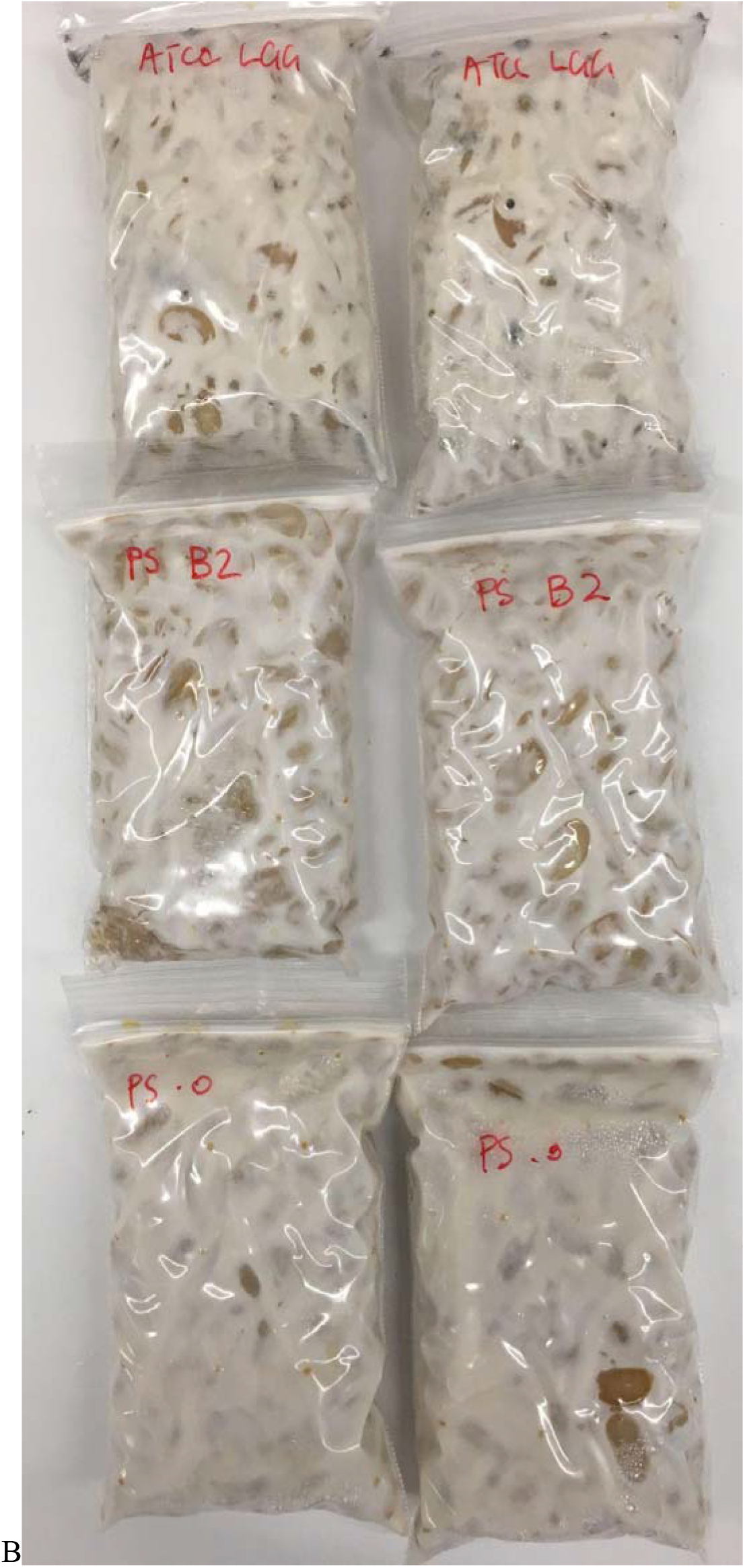
Images of tempeh made with different combinations of bacteria and fungi. A. Soybeans with no microorganisms inoculated, after incubation for 1 week. B. Soybeans with fungi and/or bacteria inoculated, after incubation for 1 week.

